# Preferential vulnerability of cortical GABAergic interneurons to *Nalcn* deficiency

**DOI:** 10.64898/2026.01.21.700772

**Authors:** Candela Barettino, Álvaro Ballesteros, Yillcer Molina, Jorge Maldonado, Mario Soriano, Leszek Lisowski, Stephan Pless, Antonio Gil-Nagel Rein, Arnaud Monteil, Ramón Reig, Isabel del Pino

## Abstract

Neuronal resting membrane potential (RMP) reflects the balance of leak conductances and varies systematically across cortical cell types. Many GABAergic interneurons exhibit more depolarized RMPs than neighbouring excitatory neurons, but the molecular mechanisms underlying these systematic differences in intrinsic excitability remain incompletely defined. The sodium leak channel non-selective (NALCN) mediates a major fraction of basal sodium conductance and cause neurodevelopmental syndromes characterized by developmental delay and cognitive impairment. Here we define NALCN contribution to cortical circuit development and function with cell-type precision. Using *Nalcn*-GFP reporter line, we map *Nalcn* expression across cortical types revealing enrichment in GABAergic hippocampal interneurons relative to hippocampal pyramidal neurons. Using conditional mouse models, we selectively deleted *Nalcn* in cortical glutamatergic lineages or forebrain GABAergic cells from early embryogenesis. Electrophysiological analysis show that developmental loss of *Nalcn* preferentially reduces intrinsic excitability across GABAergic interneuron subtypes while sparing pyramidal neurons. To understand the impact of a depolarized GABAergic RMP on brain function, we assessed the behavioral performance of *Nalcn*-deficient mice that revealed persistent deficits in contextual adaptation and spatial short-term memory. These findings reveal a neuron type-specific function of NALCN in the cerebral cortex and position NALCN as a crucial ion channel regulating basal excitability of GABAergic inhibitory circuits and cortical circuit function.

## INTRODUCTION

The electrophysiological diversity of neurons arises from the selective expression and precise subcellular allocation of ion channels^1–3^. Across cortical circuits, many GABAergic interneurons tend to exhibit more depolarized resting membrane potentials (RMP) than nearby excitatory principal neurons^4–6^, although ranges overlap across subtypes and states. While this heightened intrinsic excitability is thought to facilitate rapid inhibitory circuit recruitment shaping cortical excitability and network dynamics^7^, the molecular underpinnings of these broader RMP differences between most GABAergic interneurons and excitatory principal cells remain largely unresolved.

RMP primarily depends on K^+^ leak conductances, with additional contributions from Na^+^ leak and other resting currents (e.g. HCN/Ih). Potassium efflux is primarily mediated by members of the two-pore domain K^+^ channel family (TASK1–3) and inwardly rectifying K^+^ channels (Kir) ^8,9^. In contrast, sodium influx largely depends on a unique channel, the sodium leak channel non-selective (NALCN), which is estimated to account for up to 70% of the basal Na^+^ permeability^10,11^. NALCN function requires assembly with the ancillary subunits UNC80 (UNCoordinated 80 homologue), UNC79 (UNCoordinated 79 homologue) and FAM155A (FAMily with sequence similarity 155 member A), to form the NALCN channel complex^12^. Pathogenic loss-of-function variants in NALCN cause an ultrarare developmental encephalopathy i.e. the Infantile <u>H</u>ypotonia <u>P</u>sychomotor <u>R</u>etardation and characteristic <u>F</u>acies type <u>1</u> syndrome (IHPRF1 OMIM#: 615419). Mutations in UNC80 and UNC79 cause clinically overlapping neurodevelopmental syndromes, demonstrating that integrity of the NALCN channel complex is essential for normal brain development and function.

Functional *in vivo* studies in rodents indicate that NALCN regulates neuronal excitability in brainstem respiratory nuclei and subcortical circuits regulating REM sleep^11,13,14^. However, the perinatal lethality in global or brain-wide NALCN knockouts, the absence of NALCN-selective pharmacological tools, and limited cell-type-specific genetic interrogation have hindered the precise dissection of NALCN’s function in the cerebral cortex. Given that developmental delay and cognitive impairment are hallmark features of IHPRF1 and related syndromes, clarifying the cellular role of NALCN in cortical circuit development and function is essential to understanding how NALCN dysfunction contributes to neurodevelopmental disease.

Here, we investigate the contribution of NALCN to cortical circuit development and function, focusing on the mouse hippocampus as a well-defined model system. Using a *Nalcn-GFP* reporter mouse, we mapped *Nalcn* expression at the protein level across cortical cell types. We found that, although both glutamatergic and GABAergic neurons display *Nalcn* transcripts, NALCN is enriched in GABAergic interneurons. To test the cell-type-specific requirement for *Nalcn* during cortical development, we employed conditional mouse models in which deletion of *Nalcn* takes place in cortical glutamatergic lineages or forebrain GABAergic cells from early embryonic stages. Electrophysiological characterization revealed that developmental loss of *Nalcn* preferentially impacts the intrinsic excitability of various GABAergic interneuron subtypes while sparing glutamatergic principal neuron excitability. Behavioral phenotyping further showed that developmental disruption of *Nalcn* in GABAergic interneurons leads to abnormal adaptation to a novel context and spatial short-term memory deficits in adult mice. These findings uncover a cell-type specific requirement for NALCN in shaping cortical GABAergic interneuron excitability and inhibitory circuit function.

## RESULTS

### *Nalcn* expression is enriched in GABAergic interneurons

To understand the contribution of *Nalcn* to cortical circuit development and function, we examined the cellular distribution of NALCN/*Nalcn* and its auxiliary subunits in the cerebral cortex. We first mined published transcriptomic datasets spanning juvenile to adult mouse cortex^15^. *Nalcn* and ancillary subunits transcripts *Unc79*, *Unc80* and *Fam155A* were enriched in neuronal versus non-neuronal cell types (fig. S1A). To examine the developmental timing of *Nalcn* expression along with the expression of its auxiliary subunits across main neuronal types, we analysed a single-cell RNA sequencing dataset of the mouse cortex spanning embryonic (E10.4) to early postnatal (P) development^16^ (fig. S1B–D). Transcripts from all four NALCN complex components were detectable from late embryonic stages (E18.5) to P4 and the cell number displaying transcripts increased toward the postnatal stages in both excitatory and inhibitory neuronal populations. *Nalcn*, *Unc79* and *Unc80* showed preferential upregulation near the postnatal stages, while *Fam155A* transcription was detectable at earlier developmental timepoints (fig. S1D) in excitatory and inhibitory neurons. These data indicate that trancripts from *Nalcn* and its auxiliary subunits are present in glutamatergic and GABAergic neurons since late embryonic or early postnatal development.

To confirm NALCN protein expression, we used the *Nalcn-GFP-HA-HIS* knock-in mice^17^ in which NALCN is monitored via GFP fusion. Immunohistochemistry against GFP revealed sparse GFP signal throughout the cerebral cortex and hippocampus of knock-in animals but absent in wild type littermates. Although GFP signal was widespread, its intensity varied across neurons and layers (Fig. 1A), suggesting cell-type heterogeneity. To quantify cell-type differences between pyramidal glutamatergic and GABAergic neurons, we performed double immunofluorescence for GFP together with a maker for a major GABAergic interneuron class expressing parvalbumin (PV) (Fig. 1B). Most PV-expressing cells were GFP^+^, while PV-expressing cells constituted 20% of the GFP^+^ population (Fig 1C). Quantitative comparison of somatic GFP intensity showed that a proportion of PV^+^ interneurons expressed significantly higher GFP fluorescence than neurons allocated in the pyramidal cell layer predominantly occupied by glutamatergic neurons. To understand whether findings from PV-expressing cells generalize to other GABAergic interneurons we also performed co-stainings of GFP with somatostatin (SST) and vasoactive intestinal peptide (VIP)(fig. S2A and S2B). As seen in PV^+^ neurons, a major proportion of SST- and VIP-expressing interneurons expressed GFP (fig. S2C: 75% of SST^+^ and fig. S2D: 95% of VIP^+^), while the SST+ and VIP^+^ groups constituted a small fraction of the GFP^+^ population (fig. S2C: 17% for SST^+^ and fig. S2D: 20% for VIP^+^ cells). When GFP signal intensity was compared between SST^+^ or VIP^+^ and the pyramidal cell layer population, significant differences were found between SST^+^ and VIP^+^ subpopulations and the distribution of intensities in the pyramidal cell layer (fig. S2E and fig. S2F, respectively). Together, these data indicate that while *Nalcn* is broadly transcribed and NALCN protein is enriched in the somas of major GABAergic interneuron classes.

**Figure 1.**
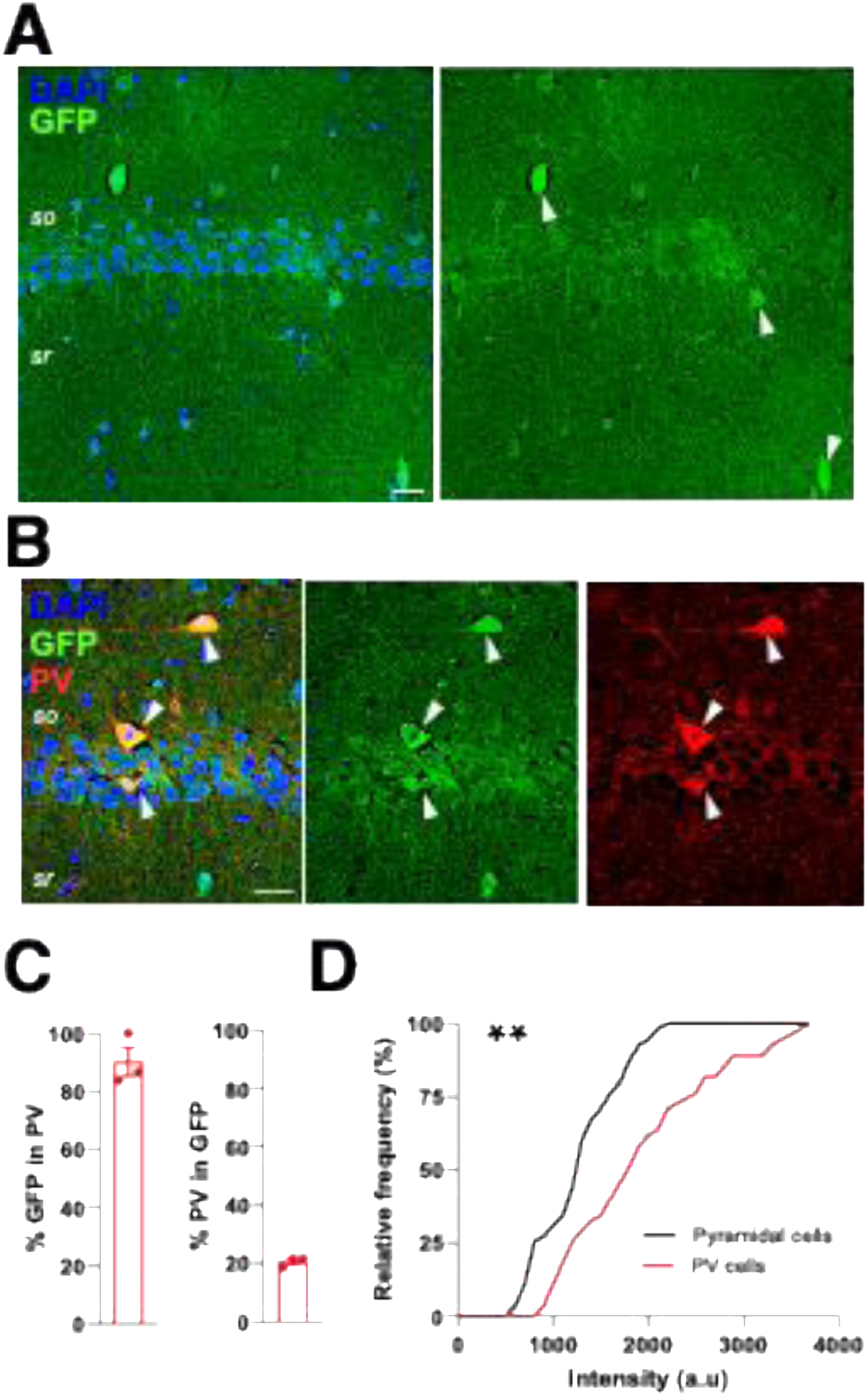
*Nalcn-GFP* expression in the hippocampal CA1. (A) Representative confocal images of the hippocampal CA1 region, showing GFP expression (green) and DAPI staining (blue). (**B**) Confocal images showing colocalization of GFP (green) with parvalbumin (PV; red) and DAPI (blue). White arrows indicate PV⁺ interneurons expressing GFP. (**C**) Percentage of GFP⁺ cells that are PV⁺ and percentage of PV⁺ cells that are GFP⁺. (**D**) Relative frequency distribution of GFP fluorescence intensity in hippocampal CA1 somas from the pyramidal cell layer (black) and PV+ somas (red) (n= 3 Nalcn-gfp-ha-his mice, ** p < 0.01, Kolmogorov-Smirnov test). Data are presented as mean ± SEM. Scale bars: 20 µm

### *Nalcn* loss attenuates the intrinsic excitability of hippocampal parvalbumin-expressing interneurons

Since *Nalcn-GFP* enrichment was observed in hippocampal GABAergic neurons, we examined whether NALCN contributes to GABAergic neuron excitability in the hippocampus through an in vivo loss of function approach. To conditionally delete *Nalcn* in forebrain GABAergic neurons from early development, we generated *Dlx5/6^(Cre/+)^;Ai9^(flox/+)^;Nalcn^(flox/flox)^*mice. First, we verified that hippocampal GABAergic neuron density and laminar distribution were comparable between *Nalcn* mutants and control *Dlx5/6^(Cre/+)^;Ai9^(flox/+)^*mice by quantifying tdTomato^+^ neuron density in CA1 subregions (fig. S3).

We next tested whether NALCN influences the electrophysiological identity of GABAergic interneurons. Whole-cell patch-clamp recordings in current-clamp mode were performed from tdTomato-labelled neurons within the *stratum oriens* of the hippocampal CA1 area (Fig. 2A). From 44 and 43 recorded cells in *Nalcn* mutants and control mice, respectively, 15 cells in *Nalcn* mutants and 16 cells in controls were confirmed *post-hoc* to express PV (Fig. 2A), showed morphologies consistent with basket and chandelier cell types (Fig. 2B) and displayed a fast-spiking firing pattern (Fig. 2C). Sholl analysis of reconstructed dendritic arbors showed a subtle but consistent increase in dendritic complexity in *Nalcn*-deficient PV^+^ neurons (fig. S4A). Unbiased clustering of intrinsic electrophysiological properties grouped control and mutant neurons into similar clusters (Fig. 2D), indicating that the lack of NALCN did not alter interneuron identity. Despite this, mutant PV^+^ cells fired substantially fewer action potentials than controls during strong depolarization steps (Fig. 2E). Analysis of action potential (AP) dynamics revealed no differences in amplitude, halfwidth, rise and decay kinetics (Fig. 2F). Analysis of passive and active electrophysiological parameters showed that a significantly more hyperpolarized resting membrane potential (RMP), increased latency to first spike at near rheobase stimulation, an enhanced medium after-hyperpolarization (mAHP) and inter-spike interval (ISI) in NALCN-deficient PV^+^ neurons when compared to control PV^+^ cells, while input resistance (Ri), rheobase and other active or passive membrane properties remained unchanged (Fig. 2G and fig S4B). These findings indicate that *Nalcn* deletion hyperpolarizes the RMP of PV^+^ interneurons and attenuates high-frequency firing resulting in reduced global excitability under strong depolarizing input.

**Figure 2.**
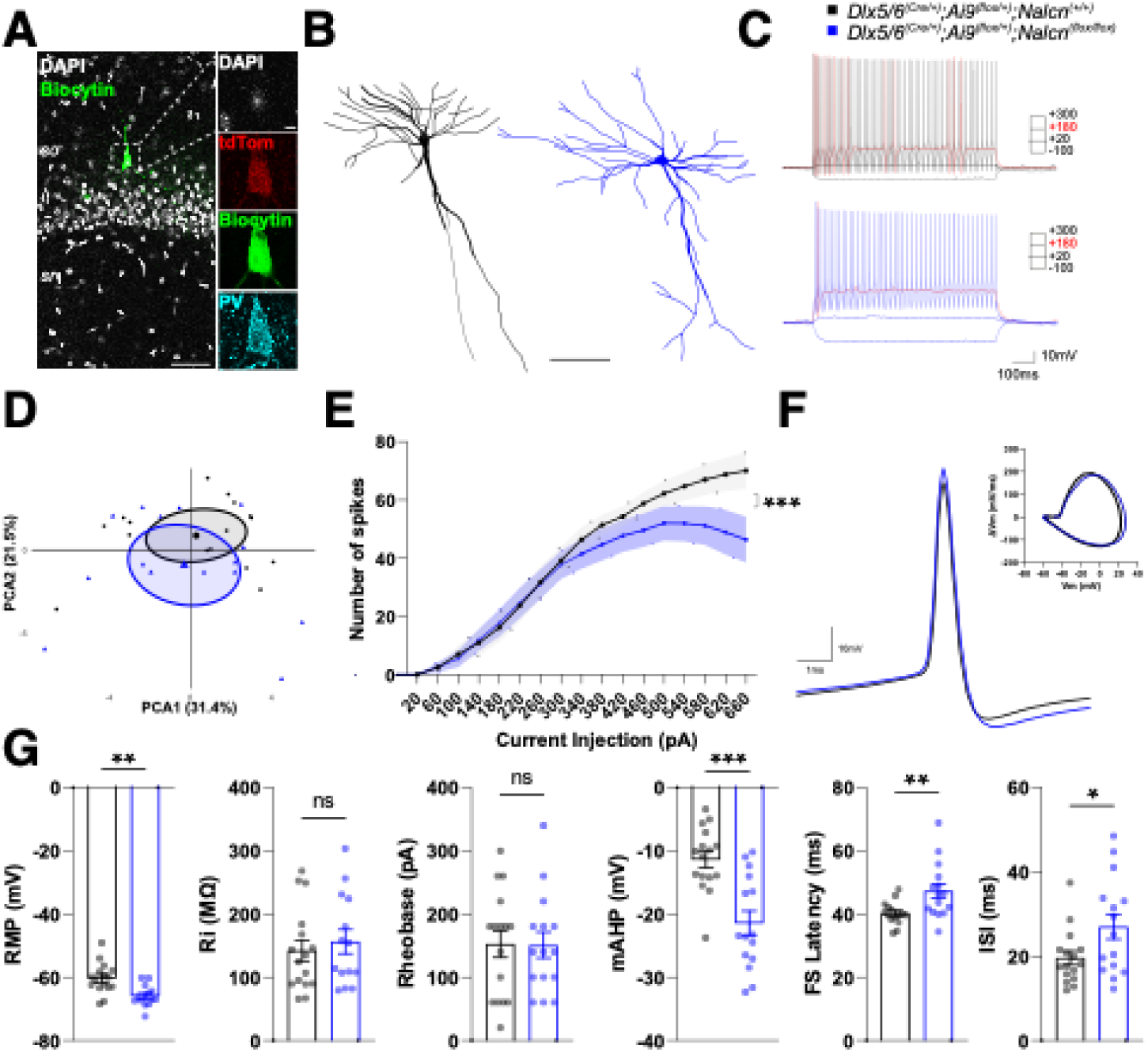
Reduced global excitability of CA1 PV^+^ interneurons in *Dlx5/6^(Cre/+)^; Ai9^(flox/+)^; Nalcn^(flox/flox)^* mice. (**A**) Representative images of biocytin-filled PV neurons (green) in CA1 *stratum oriens*. Inset: confocal image of the soma labelled with DAPI (grey), tdTomato (red), biocytin (green) and parvalbumin (cyan). (**B**) Morphological reconstructions of PV neurons from control (black) and *Nalcn* mutant (blue) mice. (**C**) Traces showing voltage responses to square pulse current injections at –100 pA, 0 pA, +180 and +300 pA. Firing at rheobase in red. (**D**) Principal component analysis (PCA) using the electrophysiological properties of PV interneurons. (**E**) Input–output relationships showing the number of action potentials evoked by incremental current injection steps from 0 to 660 pA (*** *p* < 0.001, two-way ANOVA test). (**F**) Representative trace of the first action potential (AP) from the first spike train containing ≥4Aps. Inset: phase-plane plot. (**G**) Electrophysiological parameters: resting membrane potential (RMP), Input Resistance (Ri), Rheobase, medium afterhyperpolarization (mAHP), first spike (FS) latency and inter-spike-interval (ISI) properties (ns = non-significant,* *p* < 0.05, ** *p* < 0.01; *** *p* < 0.001, unpaired two-tailed Student’s t-test). *_n =_* _16 *Dlx5/6*_*(Cre/+)_; Ai9_(flox/+)_;Nalcn_(+/+)* _and *n =* 15 *Dlx5/6*_*(Cre/+)_; Ai9_(flox/+)_; Nalcn_(flox/flox)* neurons;Data are presented as mean ± SEM. Scale bars: 50µm and 5µm in inset (A), 100 µm (B)

### *Nalcn* loss reduces the intrinsic excitability of hippocampal somatostatin-expressing interneurons

We wondered whether *Nalcn* loss could alter the intrinsic excitability of GABAergic interneurons beyond PV^+^ cells. We therefore examined *oriens-lacunosum moleculare* (O-LM) somatostatin (SST)-expressing interneurons in the hippocampal CA1 area in *Dlx5/6^(Cre/+)^;Ai9^(flox/+)^* controls and *Dlx5/6^(Cre/+)^;Ai9^(flox/+)^; Nalcn^(flox/flox)^* mutant mice (Fig. 3). SST expression and O-LM identity was confirmed by post hoc immunostaining (Fig. 3A) and morphological reconstruction (Fig. 3B).

**Figure 3.**
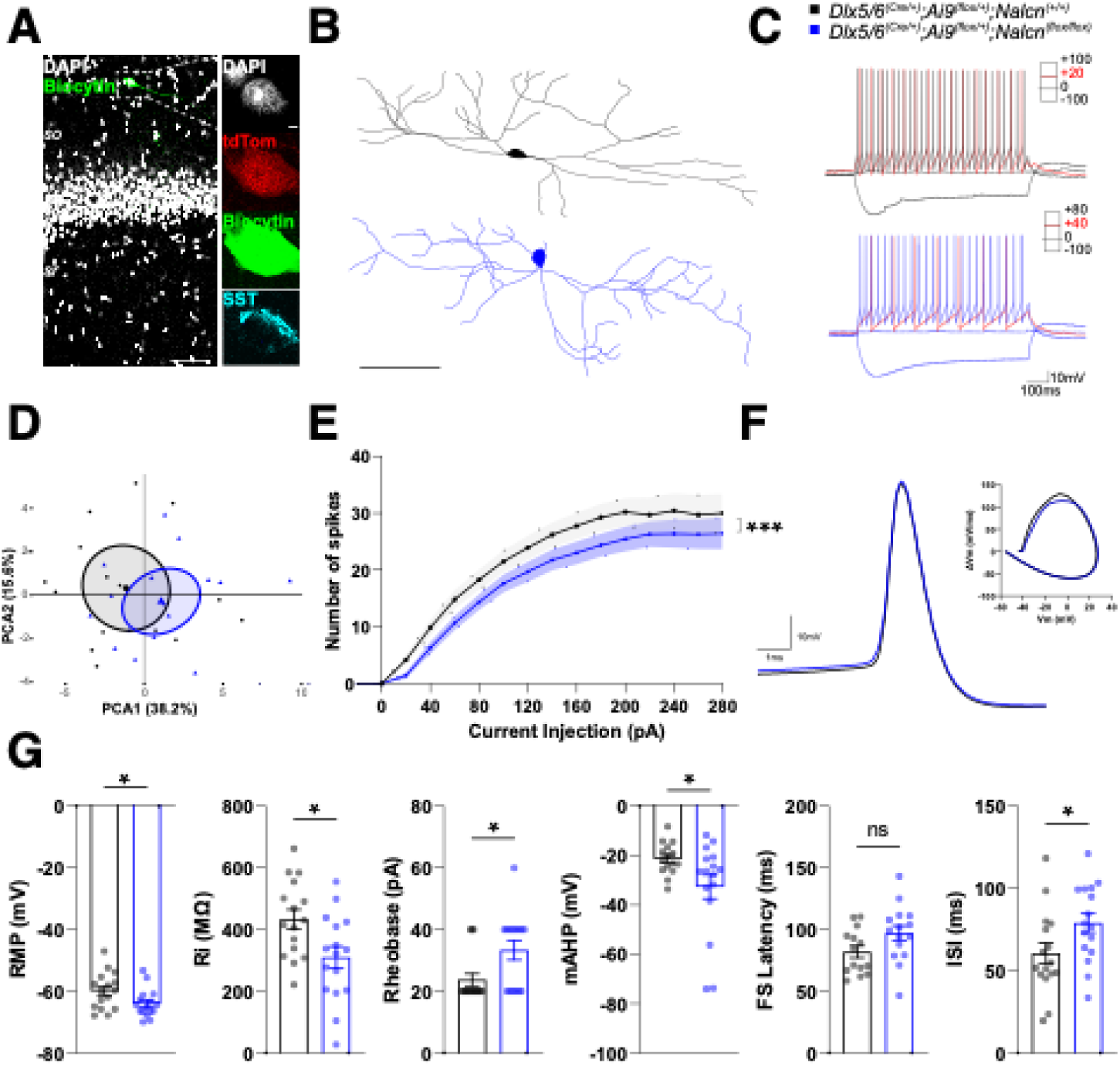
Reduced global excitability of CA1 SST^+^ O-LM interneurons in *Dlx5/6^(Cre/+)^; Ai9^(flox/+)^; Nalcn^(flox/flox)^* mice. (**A**) Representative images of biocytin-filled SST O-LM interneurons (green) located in the *stratum oriens* of hippocampal CA1. Inset: confocal image of the soma, with DAPI (grey), tdTomato (red), biocytin (green) and SST (cyan) labelling. (**B**) Morphological reconstructions of representative SST O-LM interneurons from control (black) and *Nalcn* mutant (blue) mice. (**C**) Current-clamp recordings in response to current injections (–100 pA, 0pA, +20 pA, +40 pA and +100 pA). In red, traces at rheobase for both genotypes. (**D**) Principal component analysis (PCA) plot based on electrophysiological properties of SST O-LM interneurons. (**E**) Input–output relationships showing the action potential firing as a function of increasing current injection steps from 0 to 280 pA (*** *p* < 0.001, two-way ANOVA test). (**F**) Representative trace of the first AP recorded from the first spike train containing ≥4Aps. Inset: phase-plane plot. (**G**) Intrinsic electrophysiological properties, including: resting membrane potential (RMP), Input Resistance (Ri), Rheobase, medium afterhyperpolarization (mAHP), First spike (FS) latency and inter-spike-interval (ISI) (*n =* 16 *Dlx5/6^(Cre/+)^;Ai9^(flox/+)^;Nalcn^(+/+)^* and *n =* 16 *Dlx5/6^(Cre/+)^;Ai9^(flox/+)^;Nalcn^(flox/flox)^* neurons; ns = non-significant,* *p* < 0.05, unpaired two-tailed Student’s t-test). Data are presented as mean ± SEM. Scale bars: 50µm and 5µm in inset (A), 100 µm (B)

In line with overall similarity in firing pattern observed between control and mutant SST^+^ O-LM neurons (Fig. 3C), principal component analysis of the electrophysiological parameters placed control and *Nalcn* mutant cells within a largely overlapping PCA space (Fig. 3D), indicating no categorical separation between groups. In spite of these overlapping features, analysis of input/output function revealed that mutant SST^+^ exhibit significant decrease in the action potential output across all steps of current input Fig. 3E), consistent with a global reduction in intrinsic excitability. AP waveform parameters were unchanged Fig. 3F). However, several intrinsic properties differed significantly between *Nalcn*-deficient and control SST^+^ O-LM interneurons. Specifically, *Nalcn*-deficient SST^+^ O-LM interneurons displayed a hyperpolarized RMP, reduced input resistance (R_in_), higher rheobase, enhanced mAHP and enhanced ISI (Fig. 3G and fig. S5B) when compared to control SST^+^ O-LM interneurons. These data indicate that *Nalcn* loss leads to a shift toward reduced excitability in SST^+^ O-LM interneurons.

Altogether, these results suggest that biophysical properties of multiple hippocampal GABAergic interneurons require NALCN. Loss of *Nalcn* drives a shared hypoexcitable phenotype in fast-spiking PV-expressing interneurons and in SST-expressing O-LM cells, characterized by hyperpolarized RMP, enhanced mAHP and prolonged ISI, but produces distinct GABAergic interneuron subtype-specific phenotypes. Interestingly, while PV^+^ fast-spiking interneurons primarily exhibit impaired high-frequency firing under strong depolarizing input and increase latency to first spike, SST^+^ O-LM interneurons display a broader hypoexcitable phenotype and reduced input resistance. These findings suggest that NALCN loss preserves interneuron identity in each GABAergic subtype but alters common (e.g. RMP) and distinct biophysical properties (e.g. R_in_) that sustain interneuron subtype-specific I/O function. ultimately leading to different states of hypoexcitability across GABAergic circuits.

### *Nalcn* loss does not change the global excitability of principal glutamatergic neurons

Analysis of transcriptomic datasets indicate detectable *Nalcn* expression in excitatory neuronal populations (fig.S1), raising the possibility that low-level NALCN activity may influence baseline excitability. Since low *Nalcn* expression in pyramidal neurons may have been underestimated in *Nalcn-GFP* reporter mice, we next asked whether NALCN contributes to the intrinsic properties of hippocampal pyramidal neurons. To directly test this, we selectively ablate *Nalcn* in cortical glutamatergic neurons by generating *Emx1^(Cre/+)^; Nalcn^(flox/flox)^* mice in which Cre recombinase is expressed early during pallial lineage development. Because *Emx1*-driven Cre recombination targets both cortical and hippocampal glutamatergic progenitors, we hypothesize that conditional *Nalcn* deletion could affect corticogenesis. To determine whether pallial deletion of *Nalcn* produces developmental abnormalities that could confound subsequent analysis of neuronal excitability we assessed for gross cortical morphology by measuring cortical thickness. Analysis of cortical thickness in the primary somatosensory cortex region of *Emx1^(Cre/+)^; Nalcn^(flox/flox)^* mutants and *Emx1^(+/+)^;Nalcn^flox/flox^*control mice revealed no differences (figure S6A and S6B). In addition, we addressed neuronal production and migration by quantifying density and lamination of specific glutamatergic neuron subtypes immuneractive for TBR1 and CTIP2 in the primary somatosensory cortex (S1) region of *Emx1^(Cre/+)^; Nalcn^(flox/flox)^* mutants and *Emx1^(+/+)^;Nalcn^(flox/flox)^* control mice. Neuronal density and lamination of TBR1^+^ and CTIP2^+^ glutamatergic neurons were comparable between *Nalcn* mutants and control mice (fig. S6C to S6H), indicating normal corticogenesis upon *Nalcn* loss in glutamatergic neurons.

We next performed, whole-cell recordings from deep CA1 pyramidal cells in adult *Emx1^(Cre/+)^; Nalcn^(flox/flox)^* mutants and *Emx1^(+/+)^;Nalcn^(flox/flox)^* control mice. Electrphysiological recordings were followed by morphological reconstruction and sholl analysis of dendritic morphology that revealed no morphological abnormalities in *Nalcn* deficient CA1 pyramidal neurons (Fig. 4A to C and fig. S7A). Electrophysiological analysis showed similar regular firing patterns with adaptation in both groups. Principal component analysis of intrinsic electrophysiological properties confirmed overlapping profiles between *Nalcn* mutants and controls (Fig. 4D). Furthermore input/output relationships (Fig 4E), AP dynamics (Fig. 4F) and individual electrophysiological parameters (Fig 4G and fig. S7) were unchanged. Together, these data indicate that, unlike GABAergic interneurons, developmental loss of *Nalcn* in hippocampal excitatory neurons does not affect their global intrinsic excitability.

**Figure 4.**
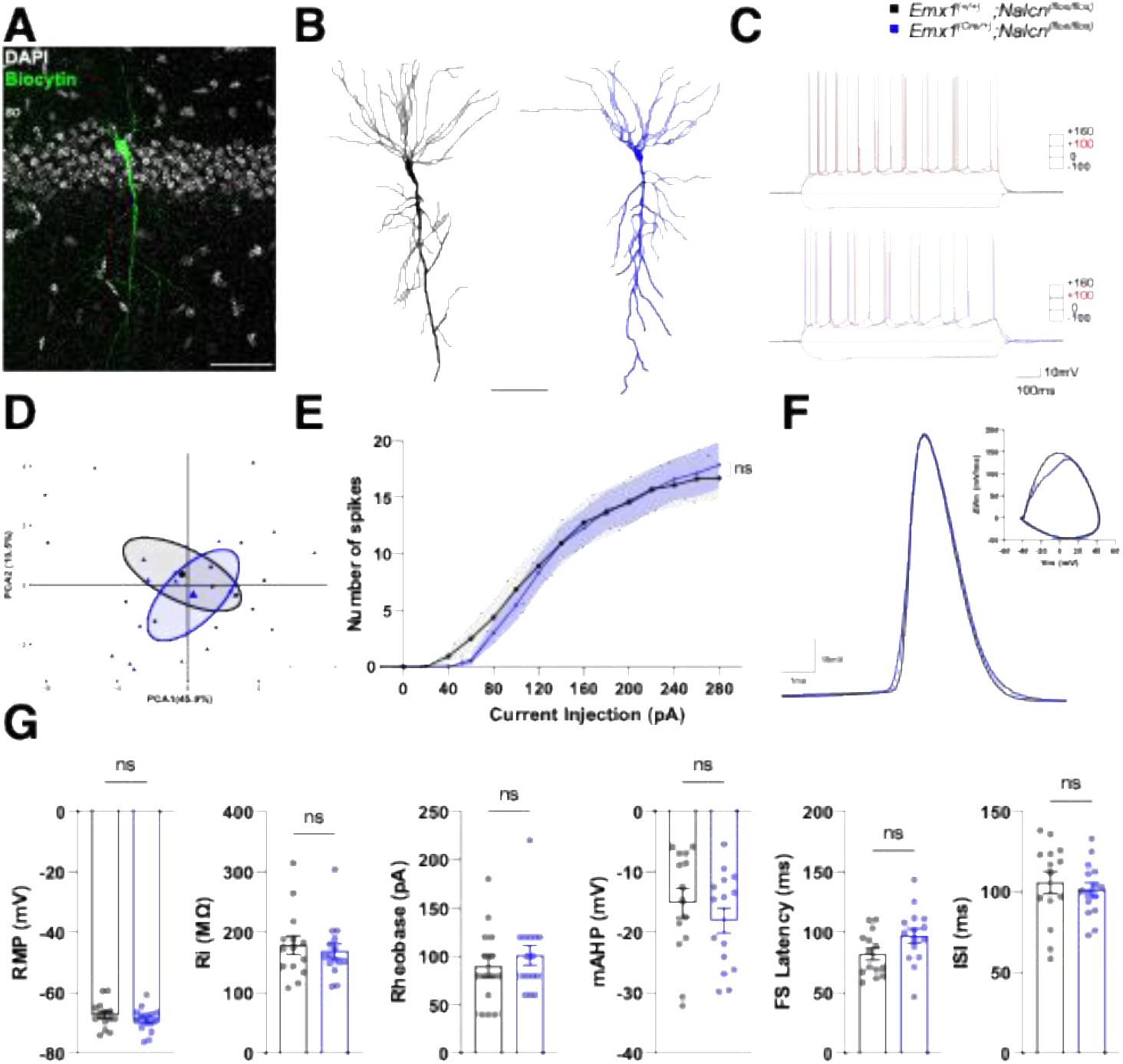
CA1 pyramidal glutamatergic excitatory neurons conserve normal electrophysiological properties upon *Nalcn* deletion in *Emx1^(Cre/+)^; Nalcn^(flox/flox)^* mice. (**A**) Representative confocal images of biocytin-filled deep pyramidal neurons (green) in the hippocampal CA1. (**B**) Morphological reconstructions of representative pyramidal neurons of *Emx1^(+/+)^;Nalcn^(flox/flox)^* (black) and *Emx1^(Cre/+)^;Nalcn^(flox/flox)^*mice (blue). (**C**) Representative traces showing voltage responses to square current pulses at −100 pA, 0 pA, +100 pA and +160 pA in deep pyramidal neurons from *Emx1^(+/+)^;Nalcn^(flox/flox)^*and *Emx1^(Cre/+)^;Nalcn^(flox/flox)^* mice. Traces at rheobase are shown in red. (**D**) Principal component analysis (PCA) using the electrophysiological properties of deep CA1 pyramidal neurons. (**E**) Input–output relationship showing action potential firing as a function of increasing current injections from 0 to 280 pA ( ns = non-significant, two-way ANOVA test). (**F**) Representative first action potential (AP) and the corresponding phase-plane plot. (**G**) Bar graphs showing resting membrane potential (RMP), Input Resistance (Ri), Rheobase, medium afterhyperpolarization (mAHP), First spike (FS) latency and inter-spike-interval (ISI) for each recorded neuron (*n =* 14 *Emx1^(+/+)^;Nalcn^(flox/flox)^*and *n = Emx1^(Cre/+)^;Nalcn^(flox/flox)^* neurons; ns = non-significant, unpaired two-tailed Student’s t-test). Data are presented as mean ± SEM. Scale bars: 50µm (A), 100 µm (B)

### *Nalcn* loss in GABAergic interneurons impairs novel context adaptation and short-term spatial memory performance

To test whether reduced global excitability observed in *Nalcn*-deficient GABAergic interneurons are relevant for brain function, we assess behavioral performance in *Nalcn* homozygous mutants (*Dlx5/6^(Cre/+)^; Nalcn^(flox/flox)^*), *Nalcn* heterozygous (*Dlx5/6^(Cre/+)^; Nalcn^(flox/+)^*) and control littermates (*Dlx5/6^(+/+)^;Nalcn^(flox/flox)^*) across a battery of assays (Fig. 5A). Locomotor activity during the first exposure to the open field was comparable between groups but mutants failed to reduce velocity and distance upon reexposure, indicating impaired habituation to a new environment (Fig5A and B). In the Y-maze, *Nalcn* deficient mice showed a similar velocity during exploration and number of total entries/alternations, but a significant reduction in the spontaneous alternation index (Fig5C), indicating a deficit in spatial working memory. In contrast, anxiety-related measures, including center-border ratio in the open field (Fig. S8A) and exploration of the elevated plus maze (Fig. 5A and D), were largely unchanged, except for a reduced preference for the light compartment in the dark-light box test (Fig5A and E). Organizational behavior measured in the nest-building test (Fig5A and fig. S8B) and depressive-like behavior measured in the forced swimming test remained unaffected (Fig5A and fig. S8C).

**Figure 5.**
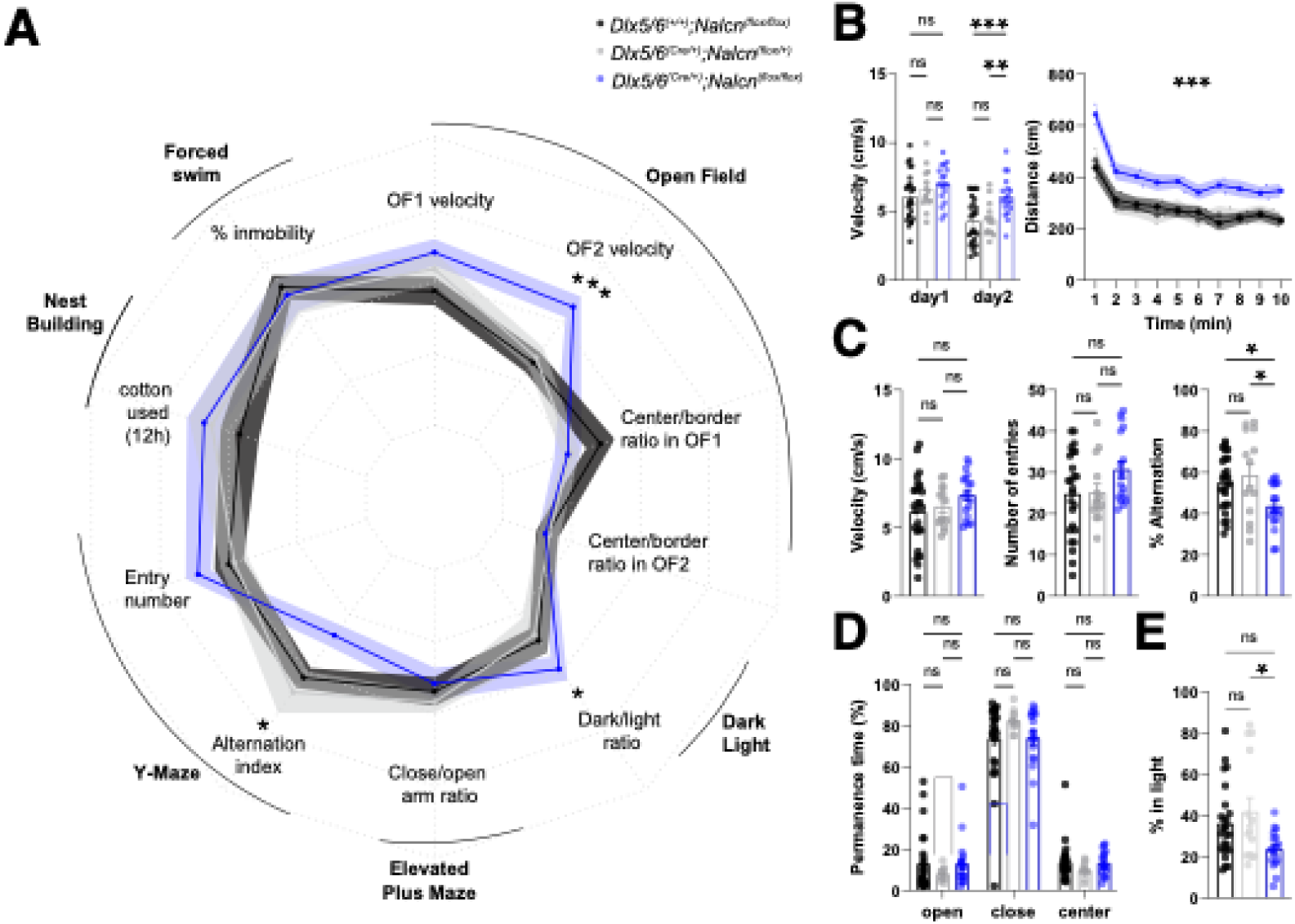
Behavioral characterisation of Dlx5/6^(Cre/+)^; Nalcn^(flox/flox)^, Dlx5/6^(Cre/+)^; Nalcn^(flox/+)^ and Dlx5/6^(+/+)^; Nalcn^(flox/flox)^mice. (**A**) Radar plot summarising the performance of *Dlx5/6^(Cre/+)^;Nalcn^(flox/flox)^*mutants (blue), *Dlx5/6^(Cre/+)^;Nalcn^(flox/+)^* heterozygous (gray) and *Dlx5/6^(+/+)^;Nalcn^(flox/flox)^* control (black) mice across diverse behavioral tasks. Parameters analysed: velocity (cm/s) on two consecutive days of open-field exploration (OF1-velocity, OF2-velocity); anxiety-like behavior measured as the centre/border exploration ratio on the two days of the open field (center/border ratio); ratio of time spent exploring dark/light boxes; ratio of time spent exploring the close/open arms in the elevated plus maze; entry number and alternation index in a spatial working memory task in the Y-maze; % of cotton used at 12h in the nesting test and percentage of time immobile in the forced swimming test. (ns = non-significant; * p < 0.05; *** p < 0.001) (**B**) Open field velocity in the two consecutive days. Total distance traveled during the second day of the open-field test, plotted in 1-min epochs (ns = non-significant, ** *p* < 0.01; *** *p* < 0.001, Two-way ANOVA with Bonferroni correction). (**C**) Velocity, entry numbers and alternation index evaluated in the Y-maze test (ns = non-significant, * p < 0.05, One-way ANOVA with Bonferroni correction). (**D**)) Percentage of time spent in closed arms, open arms, and center area in the elevated plus maze test (ns = non-significant, Two––-way ANOVA with Bonferroni correction). (**E**) Percentage of time spent in the light part of the Dark-light box test (ns = non-significant, * p < 0.05, One-way ANOVA with Bonferroni correction). (**E**) Percentage of time spent in the light part of the Dark-light box test (ns = non-significant, * p < 0.05, One-way ANOVA with Bonferroni correction). *n =* 26 *Dlx5/6^(+/+)^; Nalcn^(flox/flox)^*, *n =* 14 *Dlx5/6^(Cre/+)^; Nalcn^(flox/+)^* and *n =* 18 *Dlx5/6^(Cre/+)^; Nalcn^(flox/flox)^* mice. Data are presented as mean ± SEM.

These findings collectively indicate that conditional *Nalcn* deletion in GABAergic interneurons selectively disrupts adaptation to novelty and short-term spatial memory while leaving most affective and home cage behaviors unaffected. This suggests that NALCN-dependent excitability in inhibitory circuits is essential for adaptive response to environmental novelty and cognitive function.

## DISCUSSION

In this study, we examine the contribution of NALCN to the function of the mouse cerebral cortex. Although *Nalcn* transcripts are broadly expressed in cortical neurons, we found that NALCN protein is enriched in several GABAergic interneuron subtypes compared to glutamatergic principal cells in the hippocampus. Using conditional loss-of-function approaches, we show that deleting *Nalcn* during development in GABAergic neurons lowers their global excitability, whereas developmental deletion of *Nalcn* from pallial glutamatergic neurons does not alter their intrinsic electrophysiological properties.

At the behavioral level, mice lacking *Nalcn* in forebrain GABAergic neurons display reduced spontaneous spatial alternation and fail to habituate to a new environment, indicating deficits in cognitive performance and adaptation to environmental novelty.

Together, our findings identify NALCN as a cell-type-enriched leak conductance that sustains GABAergic interneuron excitability and is necessary for normal cognitive function and context-dependent behavioral adaptation.

### Heterogeneity in cell-type specific function of NALCN

Neurons deploy distinctive repertoires of K^+^, Na^+^ and Ca^2+^ channels that confer class-specific intrinsic firing properties and synaptic output. Although many ion channels are broadly expressed across cortical neuron types, their functional contribution is often cell type-biased. Beyond differences in expression level, this specialization can arise from neuron class-specific subcellular allocation of ion channels (e.g. HCN)^18^ or from a disproportionate functional reliance on a particular channel that has cell-type-specific redundant function, as exemplified by Cav2.1 (CACNA1A)^19^. Together, these observations support the principle that broadly expressed ion channels can exert selective functional dominance within defined neuronal populations.

In this study, we identify the sodium leak channel non-selective NALCN, previously known to mediate ubiquitous Na^+^ leak currents in cortical neurons^20,21^, as a shared molecular determinant of global intrinsic excitability across major hippocampal GABAergic interneuron classes, but not of principal glutamatergic neurons. The subclass-specific contribution of NALCN to global intrinsic excitaibility could arise from differences in subcellular localization, modulation by auxiliary subunits (UNC80^22^, UNC79^23^ and FAM155A^24^), sensitivity to extracellular Ca^2+^, or inhibition by recently identified interacting proteins (e.g. components of the SNARE complex)^21^. Thus the molecular milleux in which NALCN operates is likely a critical determinant of its functional impact across neuronal classes.

Furthermore, our findings suggest that NALCN-mediated basal sodium conductance is required to sustain a depolarized resting membrane potential (RMP) characteristic of GABAergic inhibitory interneurons, uncovering a cell-type-biased mechanism that regulates interneuron excitability. Whether principal glutamatergic neurons display a specialization in the subcellular distribution of NALCN need further investigation. Preliminary electron microscopy data suggest that NALCN localizes in dendrites within the *stratum radiatum* of the CA1 hippocampal region (data not shown), suggests NALCN contribution to principal neurons excitability could be constrained to dendritic excitability.

### Ion channels contributing to the electrophysiological diversity of cortical GABAergic interneurons

The RMP varies across neuron subtypes and reflects the weighted sum of subthreshold ionic conductances, dominated by outward K^+^ leak currents (I_L-K_) and inward Na^+^ leak (I_L-Na_). A large component of the outward K^+^ flux is mediated by members of the TASK family of two-pore domain K^+^ channels. Early work attributed RMP differences between hippocampal excitatory and inhibitory cells to TASK expression^25^, but subsequent evidence demonstrated that TASK-mediated currents are also present in multiple GABAergic interneuron subclasses, including PV^+26^ and O-LM SST^+^ cells^27^, challenging the notion that TASK channels account for interneuron-specific RMP regulation. Importantly, many of these studies relied on pharmacological tools such as the volatile anesthetic isoflurane to probe different TASK subunit contributions to I_L-K_ underpinning neuronal diversity of RMP. Because isoflurane also affects other leak channels such as NALCN^28^ and non-TASK KCNK channels^29,30^, pharmacological approaches alone are insufficient to unambiguously assign specific TASK-mediated leak currents to defined electrophysiological properties. Similarly, NALCN function has often been probed using non-specific blockers (e.g. Gd^3+^, substante P receptor antagonists, extracellular Ca^2+^). Therefore, pharmacological approaches, while indispensable, require of a new generation of ion-channel specific inhibitors to dissect the molecular underpinnings contributing to the neuron type-specific I_L-K_/ I_L-Na_ and intrinsic electrophysiological properties.

In contrast, targeted genetic deletion *in vivo* allows direct dissection of channel-specific contributions to intrinsic properties. Using this approach, we demonstrate that NALCN is necessary for establishing the RMP in hippocampal GABAergic interneurons, identifying a molecular correlate of cell-type-biased control of RMP. Our work reinforces the importance of genetic strategies for interrogating cell-specific ion channel function. In line with this idea, patch-sequencing studies demonstrate that electrophysiological properties such as resting membrane potential can not be reliably predicted from single cell transcriptomic profiles alone^31^.

These findings broaden the scope of functional specialization at the electrophysiological level to include mechanisms beyond voltage-gated conductances^19,32–34^ and suggest that inhibitory interneuron excitability also relies on discrete basal depolarizing mechanisms. Whether NALCN operates independently or in coordination with other leak channels remains an open question. Notably, compensatory interactions between NALCN and TRPC3 have been described in dopaminergic neurons^35^, raising the possibility that similar homeostatic mechanisms contribute to stabilizing excitability across cortical neuron populations.

### Translational significance of NALCN-related GABAergic vulnerability for developmental encephalopathies

Mounting evidence suggest that developmental encephalopathies converge on GABAergic interneuron dysfunction^36^. In the context of developmental channelopathies, pathogenic variants in voltage gated Na^+^, K^+^ and Ca^2+^ channels with GABAergic interneuron-biased function (e.g. SCN1A, KCNC1/Kv3.1, KCNC2/Kv3.2, CACANA1A/Cav2.1) constitute well established etiology of rare neurodevelopmental disorders^37–42^ which have been characterized at the molecular, cellular, and circuit levels^19,32,34,43,44^. In contrast, the involvement of leak conductances in inhibitory interneurons development and disease has remained poorly defined.

Mutations in genes encoding leak channels such as NALCN give rise to ultra-rare developmental encephalopathies with estimated prevalences below one in one million individuals^45–47^. Despite developmental problems and prominent cognitive impairment being a core clinical feature of IHPRF1, progress in understanding disease mechanisms has been hampered by the lethality of global or pan-neuronal *Nalcn* mutants, the lack of validated antibodies or selective pharmacological tools^13,45^. As a result, the cellular substrates linking NALCN dysfunction to cognitive and developmental phenotypes in patients have remained largely undefined.

Here we provide with the first viable mammalian model of IHPRF1 syndrome. Our model recapitulates cognitive impairments as observed in patients^48^, providing critical construct as well as face validity, and directly linking NALCN dysfunction in GABAergic neurons to cognitive outcomes. At the cellular level, our data identify GABAergic interneurons as a principal cellular target vulnerable to *Nalcn* loss, positioning IHPRF1 within the growing class of developmental interneuropathies. This GABAergic susceptibility closely parallels that observed in voltage-gated channelopathies such as Dravet Syndrome, caused by SCN1A variants, despite the fundamentally distinct biophysical role of NALCN. Thus, our work extends the concept of interneuron-driven developmental channelopathies beyond voltage-dependent conductances and highlight dysregulation of baseline excitability as a critical determinant of inhibitory circuit function and cognitive performance.

By overcoming a major experimental barrier in modelling IHPRF1 syndrome, our conditional *Nalcn* model provide a platform to dissect developmental and circuits mechanisms underlying IHPRF1 and to explore precision strategies aimed at normalizing interneuron excitability and inhibitory circuit function in developmental encephalopathies.

## MATERIALS AND METHODS

### Mice

*Nalcn-GFP-HA-HIS* knock-in line was generated in the Ren laboratory by fusing *Nalcn* to a GFP-HA-6xHis triple tag at the C-terminus^17^. Mice with conditional deletion of *Nalcn* in GABAergic neurons (*Dlx5/6^(Cre/+)^; Ai9^(flox/+)^; Nalcn^(flox/flox^*^)^) were generated by crossing *Dlx5/6-Cre* line (JAX stock #008199; RRID:IMSR_JAX:008199)^49^ with mice carrying the loxP- flanked (*flox*) *Nalcn* allele^13^ and *Ai/tdTomato* reporter line (JAX stock #007909; RRID:MGI:J:155793)^50^. Control mice carried either *Dlx5/6^(Cre/+)^; Ai9^(flox/+)^* or *Nalcn^(flox/flox^*^)^ alleles. Mice with conditional loss of *Nalcn* in pallium-derived cortical glutamatergic neurons (*Emx1^(Cre/+)^;Nalcn^(flox/flox)^*), were obtained by breeding *Emx1-Cre* lines (JAX stock #005628; RRID:IMSR_JAX:005628)^51^ with *Nalcn-floxed* mice. All animal procedures were approved by the Ethics Committee (Instituto de Neurociencias, CSIC-UMH, Spain) and complied with Spanish and European regulations for the use of laboratory animals.

### Mouse Behavioral Analysis

Behavioral testing followed standard protocols. Adult (8-12 week-old) homozygous (*Dlx5/6^(Cre/+)^; Nalcn^(flox/flox^*^)^), heterozygous (*Dlx5/6^(Cre/+)^; Nalcn^(flox/+^*^)^), and control (*Dlx5/6^(+/+)^;Nalcn^(flox/flox^*^)^) littermates were used for behavioral tests. All groups contained both male and female mice, with a balanced representation of sexes across conditions. Mice were maintained under standard housing conditions on a 12 h light/dark cycle with food and water available *ad libitum*. All tests were performed during the light phase, and the experimenter was blind to genotype. Before the behavioral assays, mice were habituated to the experimenter for 3 days. Behavioral tests were conducted in order of increasing aversiveness: open field, habituation to open field, Y-maze, elevated plus maze (EPM), dark–light box (DL-box), nest building, and forced swim test. Behavioral apparatuses were cleaned with 70% ethanol between trials and animals to avoid olfactory cues. Behavioral tasks were recorded with a Logitech C270 HD webcam and analyzed using EthoVision® XT software (Noldus, RRID:SCR_000441) unless specified otherwise.

a. Open Field (OF)

Spontaneous locomotory activity was evaluated in an open field maze, consisting of a rectangular plastic chamber (48×48×48 cm). Mice were allowed to explore the apparatus for 15 min with 70 lux constant light conditions (OF1). For analysis, the arena was divided into two regions: center (square area of 30 × 30 cm equidistant from the walls) and the remaining borders. Velocity, distance travel and the time spent in each region was calculated. Ratio center/borders was calculated to assess anxiety-like behavior according to literature. Habituation to open field was performed the day after the first exposure to the arena in the same conditions as previously described (OF2).

Spontaneous Alternation Task in the Y-maze

The spontaneous alternation task was performed in a transparent Y-maze (3 arms, 50 × 8 cm each). Mice were placed in the center and allowed to freely explore for 8 min with constant 70 lux lightning. Arm-entry sequences, total distance traveled, mean velocity, and total arm visits were quantified. Alternation index (I_alternation_) was calculated as the number of triads containing entries into all three arms divided by the maximum possible alternations as follows:

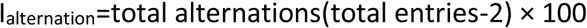

Elevated Plus Maze

The maze consisted of four arms (50 × 10 cm) two closed arms with black acrylic glass walls (30 cm high) and two open (wall-free) arms connected by a central platform, all elevated 50 cm above the floor. Light conditions were set to be 100lux in the open arms and 15 lux in the closed ones. Mice were placed in the central part of the maze and were allowed to freely explore it for 5 minutes. Time spent and the number of entries in each arm was quantified *post hoc*. Close/open arm ratio was calculated to measure anxiety-like behavior.

Dark-Light Box

Dark-Light Box maze consisted of two equal-sized compartments (25 × 25 cm), connected by a central small aperture. The light chamber was open with direct illumination of 450 lux, while the dark chamber was closed, opaque and non-illuminated. Mice were placed facing the dark side and its behavior was recorded for 5 minutes. Time spent and distance travelled in the light compartment were analysed *post hoc*, and a dark/light time ratio was calculated to assess anxiety-like behavior in an aversive light environment.

Nesting Behavior

To assess nest-building behavior, mice were housed individually and provided with two pieces of cotton fiber nesting material (3 × 1 cm each). The remaining unused material was weighed at 3, 6, 9, 12, and 24 h. The percentage of used cotton was calculated as a measure of organizational behavior as follows:

% Used cotton =weight nesting material at t 0-weight nesting material at t(x)weight nesting material at t(0)× 100

Forced Swimming Test

Mice were placed in a transparent glass cylindrical tank (12 cm diameter, 23.5 cm height) filled with water at room temperature and illuminated at 56 lux. Mice were gently introduced into the water, and their escape-related mobility was recorded for 6 min. To assess coping behavior, percentage of immobility over the total time was manually scored by an experimenter blinded to genotype.

### Analysis of Single-Cell RNA sequencing Data and Visualization

To assess the expression of *Nalcn* and its auxiliary subunits in the adult mouse brain, we analyzed the single-cell RNA sequencing dataset published by Zeisel et al., 2018^15^, that contains cells from the whole mouse nervous system, from which only brain-derived cells were selected. To evaluate the expression of *Nalcn* and its auxiliary subunits during mouse brain development, we reanalyzed the dataset from Di Bella et al., 2021^16^, containing data from the mouse cerebral cortex at multiple embryonic (E10–E18.5) and early postnatal (P1–P4) stages, covering distinct neuronal and non-neuronal subtypes.

All datasets were downloaded from the Gene Expression Omnibus (GEO, NCBI) and processed in RStudio (R version 4.0.2, RRID:SCR_001905). Analyses were carried out using the Seurat package (version 5.0.1, RRID:SCR_016341)^52^, including quality control and preprocessing steps, filtering low-quality cells (fewer than 300 detected genes) and genes expressed in fewer than three cells. Data were normalized, scaled, and variable genes identified using the SCTransform function from Seurat. Counts were log-normalized using the natural logarithm of 1 + counts per million [ln(CPM+1)]. Expression of genes across different cell types were obtained using VlnPlot function from Seurat while heatmaps were generated with pheatmap function. UMAP plots were used for dimensionality reduction visualization of developmental categories and cellular types, as well as for the expression of the evaluated genes at single-cell level.

### Immunohistochemistry

*Nalcn-GFP-HA-HIS* heterozygous adult mice (P60-P90) were used for immunohistochemistry. Mice were transcardially perfused with 4% cold PFA in 0.1M phosphate buffer (PB). Brains were post-fixed for 2h, cryoprotected in 15% and 30% sucrose in 0.1M PB, and sectioned at 40 µm thickness using a freezing microtome. For immunohistochemistry against GFP, PV, SST, VIP, CTIP2 and TBR1, slices were blocked for 2 hours at room temperature in PBS containing 0.3% Triton X-100 and 5% BSA. Overnight incubation was performed at 4°C in 0.1% Triton X-100 and 1% BSA in PBS using the following primary antibodies: chicken anti-GFP (1:500, Aves Labs Cat# GFP-1020, RRID:AB_10000240), rat anti-SST(1:100,Millipore Cat# MAB354, RRID:AB_2255365) mouse anti-PV (1:1000, Sigma-Aldrich Cat# SAB4200545, RRID:AB_2857970), rabbit anti-VIP(1:1000, Immunostar Cat# 20077, RRID:AB_572270), rabbit anti-Tbr1 (1:500, Abcam Cat# ab31940, RRID:AB_2200219) and rat anti-Ctip2 (1:500, Abcam Cat# ab18465, RRID:AB_2064130). Secondary antibodies were incubated in the same solution for 2h at RT, using the following secondary antibodies: Donkey anti-Rat Alexa-674 (1:1000, Thermo Fisher Scientific Cat#A78947, RRID:AB_2910635), Goat anti-chicken Alexa-488 (1:1000, Thermo Fisher Scientific Cat# A-11039, RRID:AB_2534096), Donkey anti-Mouse Alexa-488 (1:1000, Thermo Fisher Scientific Cat#A-21202, RRID:AB_141607), Goat anti-Rabbit biotinylated (1:1000, Vector Laboratories Cat# BA.1000, RRID:AB_2313606) and conjugated Alexa-555 Streptavidin (Thermo Fisher Scientific Cat#S32355, RRID:AB_2571525).

For biocytin-filled neurons from *ex vivo* electrophysiology experiments, 300 µm slices were fixed overnight in 4% PFA at 4°C and rinsed in PBS for three times. Slices were then blocked for 2h at RT (0.3% Triton X-100 and 5% BSA in PBS) and incubated overnight at 4°C with conjugated Alexa-488 Streptavidin (1:1000, Thermo Fisher Scientific Cat# S11223, RRID:AB_2336881) or conjugated Alexa-647 Streptavidin (1:1000, Thermo Fisher Scientific Cat# S32357, RRID:AB_2336066) in 0.1% Triton X-100 and 1% BSA in PBS. For slices in which SST and PV interneurons have been recorded, primary antibodies were added during this incubation, followed by secondary antibodies incubation using the same protocol as described before. All samples were counterstained with DAPI and mounted with Mowiol medium.

### Imaging acquisition and analysis

Confocal images were acquired using a Leica SPE II upright microscope. A 20×/0.6 NA objective was used for cell distribution analysis, and a 40×/1.15 NA objective was used for intensity measurements. A 63×/1.4 NA oil-immersion objective was employed to image the soma of Parvalbumin (PV), and Somatostatin (SST) interneurons filled with biocytin during ex vivo whole-cell patch-clamp experiments.

Image analysis was performed using FIJI (ImageJ, RRID:SCR_002285)^53^. Cell Counter plugin was used for manual quantification of colocalization between GFP and molecular markers of GABAergic interneuron subtypes (PV, SST and VIP) and density of Ctip2 Tbr1 positive neurons. Quantifications were performed across three mice, using four hippocampal slices per animal and the mean value per animal was used for statistical analysis.

For analysis of fluorescence intensity, images were acquired at 12bit depth. Intensity measurements were done by comparing the mean grey values of GABAergic interneuron somas with pyramidal-like somas within the pyramidal cell layer. Quantifications were performed across three mice, using four hippocampal slices per animal. Quantification of tdTomato-positive neuron density and laminar distribution in CA1 subregions was performed semi-automatically, and counts were normalized to total area. Cortical thickness was measured as the length (μm) of the primary somatosensory cortex (S1).

Biocytin-filled neurons from patch-clamp recordings were reconstructed using a Leica DM4B microscope equipped with a 20×/0.5 NA and Neurolucida software (MBF Bioscience; RRID:SCR_001775). Morphology analyses (Sholl analysis) were performed with Neurolucida Explorer.

### Ex vivo patch-clamp recordings and analysis

*_Dlx5/6_(Cre/+)_; Ai9_(flox/+)_; Nalcn_(+/+)* _and *Dlx5/6*_*(Cre/+)_; Ai9_(flox/+)_; Nalcn_(flox/flox)* _mice, or_ *Emx1^(+/+)^; Nalcn_(flox/flox)_* and *Emx1^(Cre/+)^; Nalcn^(flox/flox)^* mice (7–10 weeks old) were used. Mice were anesthetized with isoflurane and transcardially perfused with ice-cold artificial cerebrospinal fluid (aCSF) containing (in mM): 87 NaCl, 25 NaHCO3, 5 D-(+)-glucose, 65 sucrose, 2.5 KCl, 1.25 NaH2PO4, 0.5 CaCl2, 7 MgCl2, 5 ascorbic acid, and 3.1 pyruvic acid, saturated with 95% O2 / 5% CO2 (pH 7.4). Brains were rapidly removed and placed in ice-cold oxygenated aCSF. Sagittal slices (300 µm) were prepared using a vibratome (Leica VT-1000) and transferred to a holding chamber containing aCSF composed of (in mM): 125 NaCl, 25 NaHCO3, 25 D-(+)-glucose, 2.5 KCl, 1.25 NaH2PO4, 2 CaCl2, 1 MgCl2, 1 ascorbic acid, and 4 pyruvic acid, bubbled with 95% O2 / 5% CO2 (pH 7.4). Slices were allowed to recover at 35 °C for 30 min.

For patch clamp recordings in whole-cell configuration, slices were transferred to a chamber and continuously superfused with aCSF. Recordings were performed using a potassium gluconate–based internal solution containing (in mM): 135 K-gluconate, 10 HEPES, 10 Na-phosphocreatine, 4 KCl, 4 MgATP, and 0.3 NaGTP, adjusted to 290 mOsm and pH 7.2–7.4. Biocytin (1.5–2.5 mg/mL) was included in the internal solution for post-hoc immunohistochemistry. tdTomato-expressing neurons were visualized with an upright microscope (LNscope 240XY; Luigs & Neuman) equipped with an ORCA-Spark digital CMOS camera (C11440-36U, Hamamatsu), a 40×/0.8 NA water-immersion objective (Olympus), a pE-300white fluorescence lamp (CoolLED; RRID:SCR_021073), and infrared-differential interference contrast optics. Patch pipettes (6–10 MΩ) were pulled from borosilicate glass using either a vertical puller (PC-100, Narishige) or a horizontal puller (P-1000, Sutter Instrument, RRID:SCR_021042). Electrophysiological data were acquired at 20 kHz using a MultiClamp 700B amplifier (Molecular Devices, RRID:SCR_018455), a DigiData 1550B digitizer (Molecular Devices), and pClamp software (Molecular Devices, RRID:SCR_011323). Only cells in which access resistance (R_a_) was <25MΩ upon break in and ΔR_a_ < 20% during the course of the experiment were included in the analysis.

Current-clamp protocols were used to obtain intrinsic electrophysiological properties. Resting membrane potential (RMP) was measured during 1 min of a gap-free protocol after cell membrane break-in. Input resistance (Rin) was calculated using 800 ms hyperpolarizing steps (-40 to +20 pA, Δ20 pA). Rheobase, action potential properties, and input–output curves were evaluated using 800 ms depolarizing steps (-100 to +280 pA, Δ20 pA) from RMP. For fast-spiking PV interneurons, the mentioned parameters were obtained using 800 ms depolarizing steps (-100 to +660 pA, Δ40 pA) from RMP. Threshold potential for spikes was defined as dV/dt = 10 mV/ms, using the first derivative method, and AP threshold was defined in a region of 10ms before peak. Rheobase was defined as the minimum current injected to obtain the first AP from neurons held at-70mV. AP rise and decay time were determined in a minimum-maximum cutoff percentage of 10-90%. Half-width is calculated as the time between rise and decay at 50%amplitude. fAHP and mAHP were detected in a search region of 0–9 ms and 30–70 ms post-peak, respectively. Inter-spike interval (ISI) was defined as the interval between the first and second AP at the first current step with ≥4 spikes. Spike frequency adaptation (SFA) was obtained by the divisor method, dividing the first ISI by the final ISI of the trace with ≥4AP. Phase-plane plots were generated by plotting the first derivative of the membrane potential (dV/dt) against the membrane voltage (Vm) of the first spike of the trace with ≥4AP. Data analysis was performed offline with EasyElectrophysiology software v2.7.2 (http://www.easyelectrophysiology.com/) RRID:SCR_021190).

## Statistical analysis

Statistical analyses were carried out with the GraphPad Prism 9 software (RRID:SCR_002798) and with R programming. Principal component analysis was performed in R. All data are presented as mean ± SEM. Statistical methods were used to predetermine sample sizes. Randomization was not applied. Experiments and analyses were performed blind to genotype. Biological replicates (n values) represent different animals for behavior and immunohistochemistry, or cells from more than three different brains for electrophysiology, sourced from more than three different litters. Differences were considered significant when p < 0.05 (*), p < 0.01 (**) or p < 0.001 (***). Prior to statistical comparisons, all datasets were tested for normality (Shapiro–Wilk test) and for homoscedasticity (Levene’s test). Parametric tests, including unpaired two-tailed Student’s t-test, one-way ANOVA or two-way ANOVA, were applied when data met both normality and homoscedasticity assumptions. For datasets that did not meet these criteria, non-parametric tests, such as the Mann–Whitney U test, were used. Multiple comparisons were corrected using the Bonferroni method. Both male and female animals were included in all experiments to account for potential sex differences.

## ACKNOWLEDGEMENTS

We are grateful to Prof. Dejian Ren (University of Pennsylvania) and Prof. Stephan Pless (University of Copenhagen) for insightful discussions and ideas. We also thank Prof. Dejian Ren for kindly providing the *Nalcn-GFP-HA-HIS* and *Nalcn-flox* mouse lines. We thank Alexandra Typou, Alessandra Miatton, Mónica Peralta, Victoria Ramos and Alicia Juan for technical assistance, to Isabel Pastor from the Fundación Libellas Foundation and to members of the Channeling Hope Foundation and members of the Del Pino, Reig, Flames and Leyva laboratories for stimulating discussions.

## Author contributions

Conceptualization, I.D.P; Formal Analysis, C.B., A.B.G., Y.M., J.M., I.D.P.; Investigation, C.B., A.B.G., Y.M., J.M., M.S., L.L., S.P., A.G-N.R., A.M., R.R., I.D.P.; Resources, L.L., A.G-N.R., S.P., A.M., R.R., I.D.P., Writing – Original Draft, C.B., A.B.G, I.D.P; Writing – Review and Editing, C.B., A.B.G., Y.M., J.M., L.L., S.P., A.G-N.R., A.M., R.R., I.D.P.; Visualization, C.B., A.B.G., R.R., I.D.P.; Supervision, R.R., I.D.P.; Project Administration, R.R., I.D.P

## Data and code availability

All data needed to evaluate the conclusions in the paper are present in the paper and/or in the Supplementary Materials.

